# Identification of human tendon cell populations in healthy and diseased tissue using combined single cell transcriptomics and proteomics

**DOI:** 10.1101/2019.12.09.869933

**Authors:** AR Kendal, T Layton, H Al-Mossawi, R Brown, C Loizou, M Rogers, M Sharp, S Dakin, L Appleton, A Carr

## Abstract

The long-term morbidity of tendon disease in an increasingly active and ageing population represents a growing area of unmet clinical need. Tendon disorders commonly affect the lower limb, ranging from isolated tendon rupture to degenerative joint deformity. In the absence of valid animal models of chronic tendinopathy, traditional methods to isolate and identify crucial sub types involved in disease are limited by the heterogeneity of tendon cells, by their relative paucity in tissue and by the density of the surrounding collagen matrix. To overcome this, we have used next generation CITE-sequencing to combine surface proteomics with in-depth, unbiased gene expression analysis of single cells derived ex vivo from healthy and diseased tendon.

For the first time we have been able to show that human tendon consists of at least eight sub-populations of cells. In addition to endothelial cells, Tc cells, and macrophages, there are five distinct tenocyte populations expressing *COL1A* genes. These consist of a population of resident cells expressing microfibril associated genes (*FBN1, VCAN*, *DCN, EMILIN1*, *MFAP5*), a group of *SCX*+ cells co-expressing high levels of pro-inflammatory markers, a population of *APOD+* fibro-adipogenic progenitors (FAPs), *TPPP3/PRG4+* chondrogenic cells *(COMP, CILP, PRG4)* and *ITGA7*+ Smooth Muscle-Mesenchymal Cells, recently described in mouse muscle but not, as yet, in human tendon. Surface proteomic analysis identified markers by which these sub-classes could be isolated and targeted in future.

In comparison to healthy tendon, diseased tendon harboured a greater proportion of *SCX+* tendon cells and these expressed high levels of pro-inflammatory markers including *CXCL1, CXCL6, CXCL8, PDPN* and previously undescribed *PTX3. W*e were also able to show that whereas disease associated genes such as *CD248* and *PDPN* were expressed by *COL1*+ tenocytes, *IL33* was restricted to endothelial cells of chronically diseased tendon.

## Introduction

Musculoskeletal disorders are responsible for the second largest number of years lived with disability worldwide(1). The morbidity associated with tendon degeneration in an ageing, and increasingly active, population represents an escalating challenge to healthcare services. Tendinopathy affects up to third of the population, accounting for 30% of primary care consultations (2–4). Tendon disorders commonly affect the lower limb and result in long term pain and disability. These range from isolated tendon rupture (most commonly the Achilles tendon), to disease that drives complex foot deformity, such as adult acquired flat foot deformity (AAFD) affecting 2-3% of the adult population (5–7).

Early research into patients with Marfan syndrome demonstrated that tendon cells are surrounded by peri-cellular matrix microfibrils formed by fibrillin chains and bound ancillary proteins (including versican, fibulin, matrix associated glycoproteins). They represent a mechanism for altering growth factor signalling (e.g. sequestering TGF-beta), controlling morphogenic gradients, and influencing cell interactions with the extracellular collagen matrix (8–10). They may allow resident tenocytes to sample, respond and influence tendon structure and function (11). The development of new therapies involves understanding how key sub-populations of tendon cells/fibroblasts are able to maintain a dense tendon extracellular matrix and importantly, how they survey and respond to events which threaten tissue integrity.

The bulk of tendon consists of dense collagen extracellular matrix and residing cells tend to be sparse (particularly in healthy tendon), heterogenous and auto-flourescent; so limiting previous techniques (e.g. flow cytometry) for studying cells of interest. Moreover the mechano-sensitivity of tendon cells risks confounding their interrogation in vitro including the associated changes in gene expression (12). It is currently not possible to define, isolate and target specific sub-populations of matrix producing tendon cells involved in human disease. Single cell RNA sequencing offers an unbiased, agnostic and sensitive inventory of the transcriptome of individual cells and allows characteristics of groups of cells based on shared and differential gene expression data (13). This approach has been successfully used to characterise cell types is mouse tissue (14–17).

CITE-Seq (Cellular Indexing of Transcriptomes and Epitopes by Sequencing) is a novel iteration of single cell sequencing that uses oligonucleotide barcodes conjugated to monoclonal antibodies to combine surface proteomics with single cell RNA/transcriptomic information. To our knowledge this is the first time CITE-Seq has been applied to healthy and diseased human tendon. We have identified multiple sub-populations of cells in human tendon, five of which show increased expression of *COL1A* genes. These include two groups that co-express microfibril genes, a group expressing genes associated with fibro-adipogenic progenitors (FAPs), a *TPPP3/PRG4*+ chondrogenic group and *ITGA7*+ Smooth Muscle-Mesenchymal Cells (SMMCs), previously described in mouse but not in human tendon(16, 17). These findings support the presence of multiple specialised tendon cell sub-types and open new avenues to interrogate key cell pathways that underpin chronic tendon disease.

## Methods

### Collection of tissue tendon samples donated by patients

Tendon biopsies were collected from patients with informed donor consent under ethics from the Oxford Musculoskeletal Biobank (09/H0606/11) in compliance with National and Institutional ethical requirements. Only waste tissue that would otherwise have been disposed of was collected. Healthy tendon samples were obtained form patients undergoing tendon transfer procedures. These included patients undergoing hamstring (gracilis and semimembranosus) tendon reconstruction of knee anterior cruciate ligament; reconstruction of irreparable Achilles tendon rupture using flexor hallucis longus tendon; and tibialis posterior tendon transfer procedures to improve foot biomechanics. Diseased tendon samples were obtained from patients undergoing surgical exploration and debridement of intractable, chronic painful tendon disease. These included tendinopathic peroneus longus, the Achilles tendon and the extensor digitorum tendon associated with a painful fixed flexed deformity of the proximal inter-phalangeal joint (‘Hammer toe’).

### Tendon sample digestion

Samples were immediately placed in 4°C Iscove’s Modified Dulbecco’s Medium (IMDM) without antibiotics and without FCS. The tendons were rinsed in 1 x PBS, cut axially using a size 10 surgical scalpel into 1mm^3^ pieces and incubated at 37°C for 45 mins in LiberaseTM (Merck) and 10ul/ml DNAse I (Thermo Scientific™). Ham’s F-12 media + 10% FCS was added and the digested tissue passed through a 100µm cell strainer. The cells suspension was centrifuged at 350g for 5 mins and re-suspended in 1ml of FCS:DMSO (9:1) freezing medium and immediately placed in −80°C storage for future batch analysis.

### CITE-seq

Digested healthy and diseased samples were defrosted, washed and re-suspended in 100µl staining buffer (1 × PBS + 2% BSA + 0.01% Tween). As per the CITE-Seq protocol (https://citeseq.files.wordpress.com/2019/02/cite-seq_and_hashing_protocol_190213.pdf), cells were then incubated for 10 minutes at 4°C in human Fc Blocking reagent (FcX, BioLegend). Cells were incubated at 4°C for a further 30 minutes with 0.5µg of TotalSeq™-A (Biolegend) monoclonal anti-CD10, anti-CD105, anti-CD146, anti-CD26, anti-CD31, anti-CD34, anti-CD44, anti-CD45, anti-CD54, anti-CD55, anti-CD90, anti-CD95, anti-CD73, anti-CD9 and anti-CD140a antibodies (see Table S1). In addition each sample cell suspension was incubated with 0.5µg of the relevant cell hashing antibodies (Biolegend). The cells were then washed three times with staining buffer and re-suspended in 1 × PBS at 1000 cells/µl. The cells suspensions were filtered using a 100µm sieve. The final concentration, single cellularity and viability of the samples, was confirmed using a haemocytometer. Cells were loaded into the Chromium controller (10x-Genomics) chip following the standard protocol for the Chromium single cell 3’ kit. A combined hashed cell concentration was used to obtain an expected number of captured cells between 5000-10000 cells. All subsequent steps were performed based on the CITE-Seq protocol (https://citeseq.files.wordpress.com/2019/02/cite-seq_and_hashing_protocol_190213.pdf). Libraries were pooled and sequenced across multiple Illumina HiSeq 4000 lanes to obtain a read depth of approximately 30,000 reads per cell for gene expression libraries.

The raw single-cell sequencing data was mapped and quantified with the 10x Genomics Inc. software package CellRanger (v2.1) and the GRCh38 reference genome. Using the table of unique molecular identifiers produced by Cell Ranger, we identified droplets that contained cells using the call of functional droplets generated by Cell Ranger. After cell containing droplets were identified, gene expression matrices were first filtered to remove cells with > 5% mitochondrial genes, < 200 or > 5000 genes, and > 25000 UMI. Downstream analysis of Cellranger matrices was carried out using R (3.6.0) and the Seurat package (v 3.0.2, satijalab.org/seurat).

After quality control filtering, data were normalised for RNA gene expression, hashed antibody (HTO) and surface antibody (ADT) expression level. Based on the HTO expression level, we were able to subsample cells from a particular donor tendon (Figure S2). Normalised data from all healthy tendon cells were combined into one object and integrated with data from cells of diseased tendon. Variable genes were discovered using the SCtransform function with default parameters. The FindIntegrationAnchors function command used default parameters (dims = 1:30) to discover integration anchors across all samples. The IntegrateData function was run on the anchorset with default additional arguments. ScaleData and RunPCA were then performed on the integrated assay to compute 15 principal components (PC). Uniform Manifold Approximation and Projection (UMAP) dimensionality reduction was carried out and Shared Nearest Neighbour (SNN) graph constructed using dimensions 1:15 as input features and default PCA reduction (18). Clustering was performed on the Integrated assay at a resolution of 0.5 with otherwise default parameters which yielded a total of 12 clusters, each composed of cells originating from both healthy and diseased samples across the study patients (19).

### Histology

Immunohistochemistry was performed on a Leica Bond™ system using the standard protocol F30. The sections were pre-treated using heat mediated antigen retrieval with citrate-based buffer (pH6, epitope retrieval solution 1) or EDTA based buffer (pH9, epitope retrieval solution 2) for 20 mins. The sections were then incubated with antibody for 30 mins at room temperature and detected using an HRP conjugated polymer system in which DAB was used as the chromogen. The sections were counter-stained with haematoxylin and mounted with Aquatex. The following antibodies were used; anti-Pentraxin 3/PTX3 [MNB1] (Abcam 90806), anti-Oligodendrocyte Specific Protein (Abcam 53041), anti-Cytokeratin 7 [RCK105] (Abcam 9021), anti-ITGA7 (Abcam 203254), anti-MGP (Abcam 86233), anti-TEM1 (Abcam 67273), anti-CD31 [JC70A] (DAKO 20057487), anti-CD68 (DAKO 20058607), anti-Smooth muscle actin (Abcam 5694), anti-periostin [EPR20806] (Abcam 227049), anti-APOD (orb155698), anti-CXCL14 (Abcam 137541), anti-Podoplanin [D2-40] GTX31231 (lot 821903108), anti-CLDN11 (Abcam 53041), anti-Cysteine Dioxygenase CDO1 (Abcam 234693).

## Results

### A single cell atlas of human tendon in health and disease

In order to characterise healthy and diseased human tendon cell subtypes, we performed scRNA-seq and CITE-seq integrated analysis of cells from eight healthy and eleven diseased tendon samples. Cells were incubated with one of eight surface hashing antibodies so that their sample of origin could be identified following sequencing in one of three lanes. After normalization and quality control of the transcriptome data, cells were selected based on high ‘expression level’ of their donor specific surface hashing antibody and low ‘expression level’ of the remaining hashing antibodies. In total, 11,948 cells with an average of 1,700 genes per cell were selected for ongoing analysis. Of these single cells, 6,411 were obtained immediately ex-vivo from digested healthy and diseased tendon and all subsequent data is derived from this group of cells. The remainder were cells from P1 culture of two healthy and one diseased tendon to allow comparison with ex-vivo single cell sequencing (Figure 1, S4, see Discussion).

**Figure 1.**
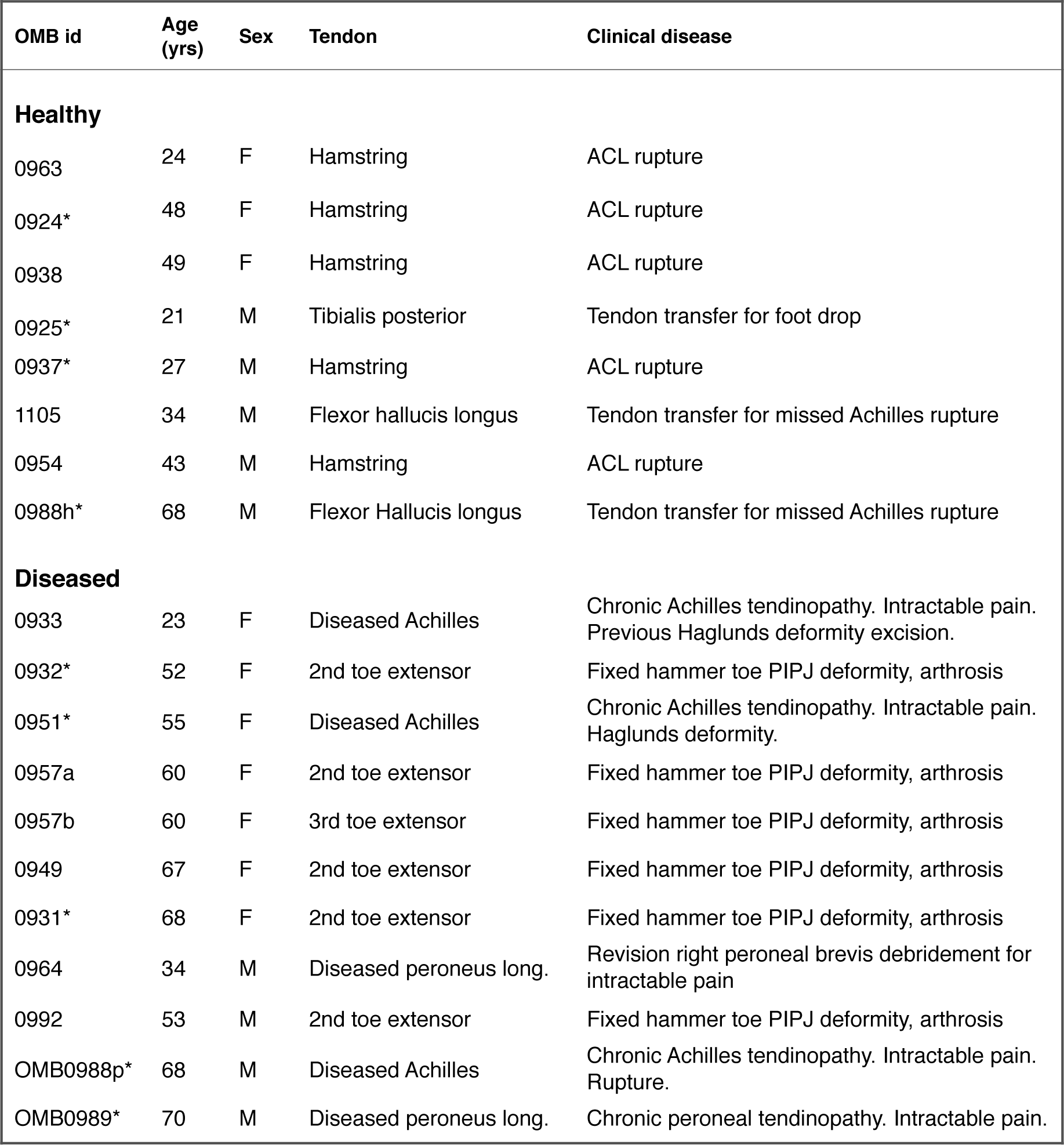
Patient donor demographics of healthy and disease tendon. Healthy tendon was obtained from patients who underwent tendon transfer procedures for reconstruction of knee anterior cruciate ligament (ACL), or ruptured Achilles tendon or to treat foot drop. Diseased tendon samples were restricted to patients who had chronic tendinopathy and medically intractable pain. Diseased tendon samples were from significantly older patients than healthy tendon (mean age 55 years v 39 years, p= 0.04). * Used for immuno-histochemistry.

Unsupervised graph based clustering was performed on the integrated dataset of both diseased and healthy cells obtained ex-vivo and was visualized using UMAP (uniform manifold approximation and projection)(18). This resolved eight distinct transcriptomic clusters found in both healthy and diseased tendon (Figure 2). Consistent with previous observations, a greater number of cells were obtained from diseased compared to healthy tendon (Figure 2A). Each cluster was further annotated by combining the top differentially expressed genes with a set of literature-defined gene markers and revealed five *COL1A* expressing tenocyte clusters (initially labelled ‘Tenocyte A-E’), a combined group of endothelial cells, macrophages and Tc lymphocytes (Figure 2A,B).

**Figure 2.**
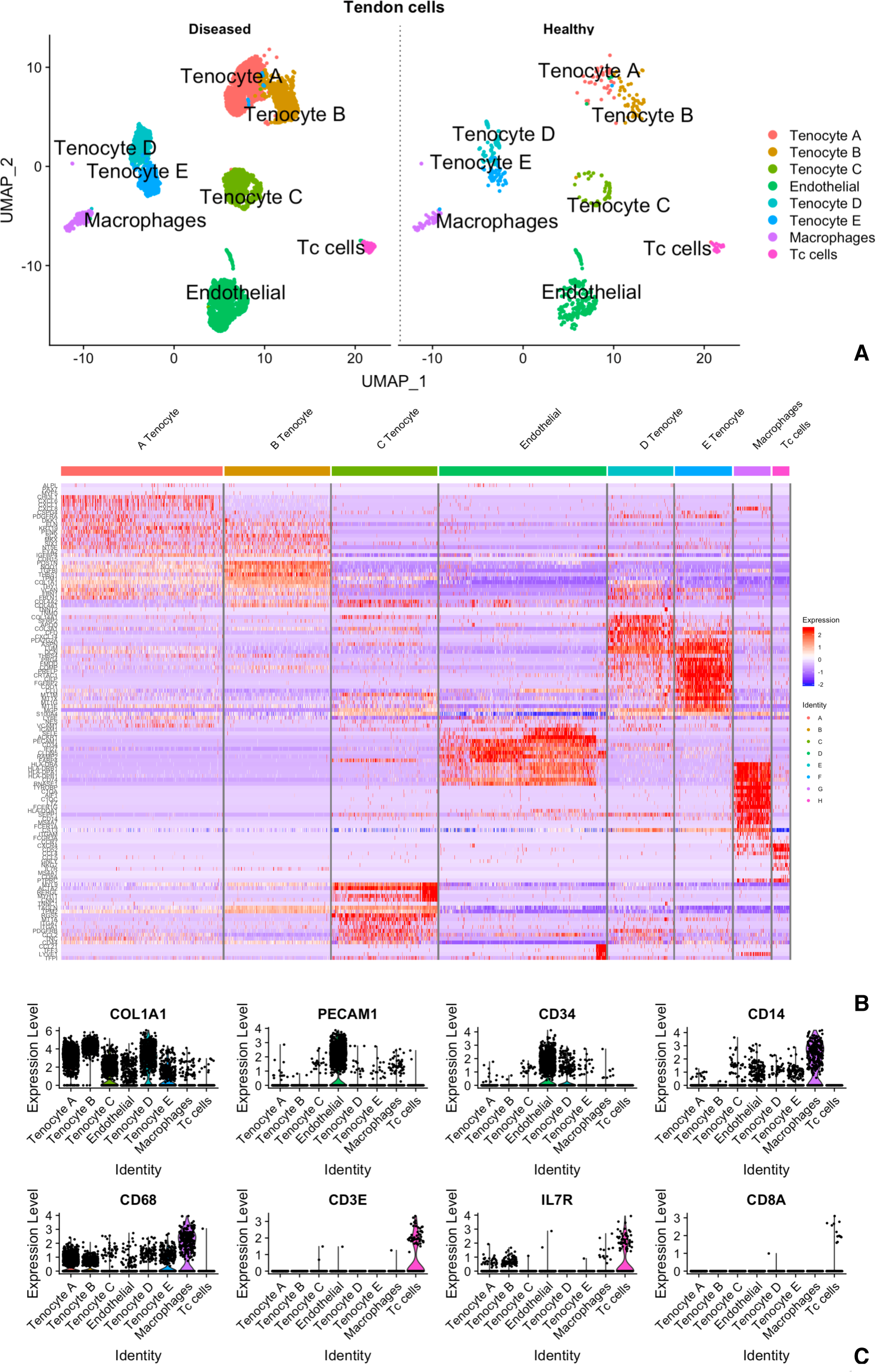
A single cell gene atlas of human tendon in health and disease. **(A)** Uniform Manifold Approximation and Projection (UMAP) dimensionality reduction was performed and Shared Nearest Neighbour (SNN) graph constructed using dimensions 1:15 as input features with default PCA reduction. Eight distinct cell populations were found on clustering of the Integrated assay at a resolution of 0.5 with otherwise default parameters. Each cluster composed of cells originating from both healthy and diseased samples across the study patients. **(B)** RNA expression heatmap for clusters (coloured columns) and genes (rows). Genes were chosen based on unbiased analysis of the top 50 differentially expressed genes, the top 4 genes per cluster and literature selected markers. **(C)** Violin plots summarising the expression pattern of selected markers to identify each major cluster.

### Multiple distinct tenocyte populations reside in human tendon

The five cell clusters that expressed tendon matrix *COL1A1/2* were provisionally labelled Tenocyte A-E. In order to delineate these groups further, the expression of genes coding for the commonest matrix proteins found in human tendon (20) was compared across the five clusters (Figure 3A, dot plot). The gene expression of any given *COL1A+* cluster was also directly compared to the remaining four clusters to highlight any additional differences between clusters (Figure 3B, volcano plots). Together these demonstrated that Tenocyte A and Tenocyte B clusters contained cells expressing *SCX* as well as genes associated with extracellular tendon microfibrils (*FBN1, MFAP5, VCAN, EMILIN1*), and TGFβ signalling (e.g. *TGFB1*, *LTBP1*, *LTBP2*). In order to discriminated between these two clusters, their gene expression profiles were compared directly to each other (Figure 3B, final volcano plot). Tenocyte A cells displayed up-regulation of *KRT19, CHI3L, CLDN11, BAALC, PENK, SERPINE2* and pro-inflammatory genes *CXCL1, CXCL6, CXCL8, PTX3*. In comparison, Tenocyte B cells expressed higher levels of *COL4A1, KRT7, POSTN, TAGLN, THBS1, TIMP3, IGFBP5/7* and *LTBP2*.

**Figure 3.**
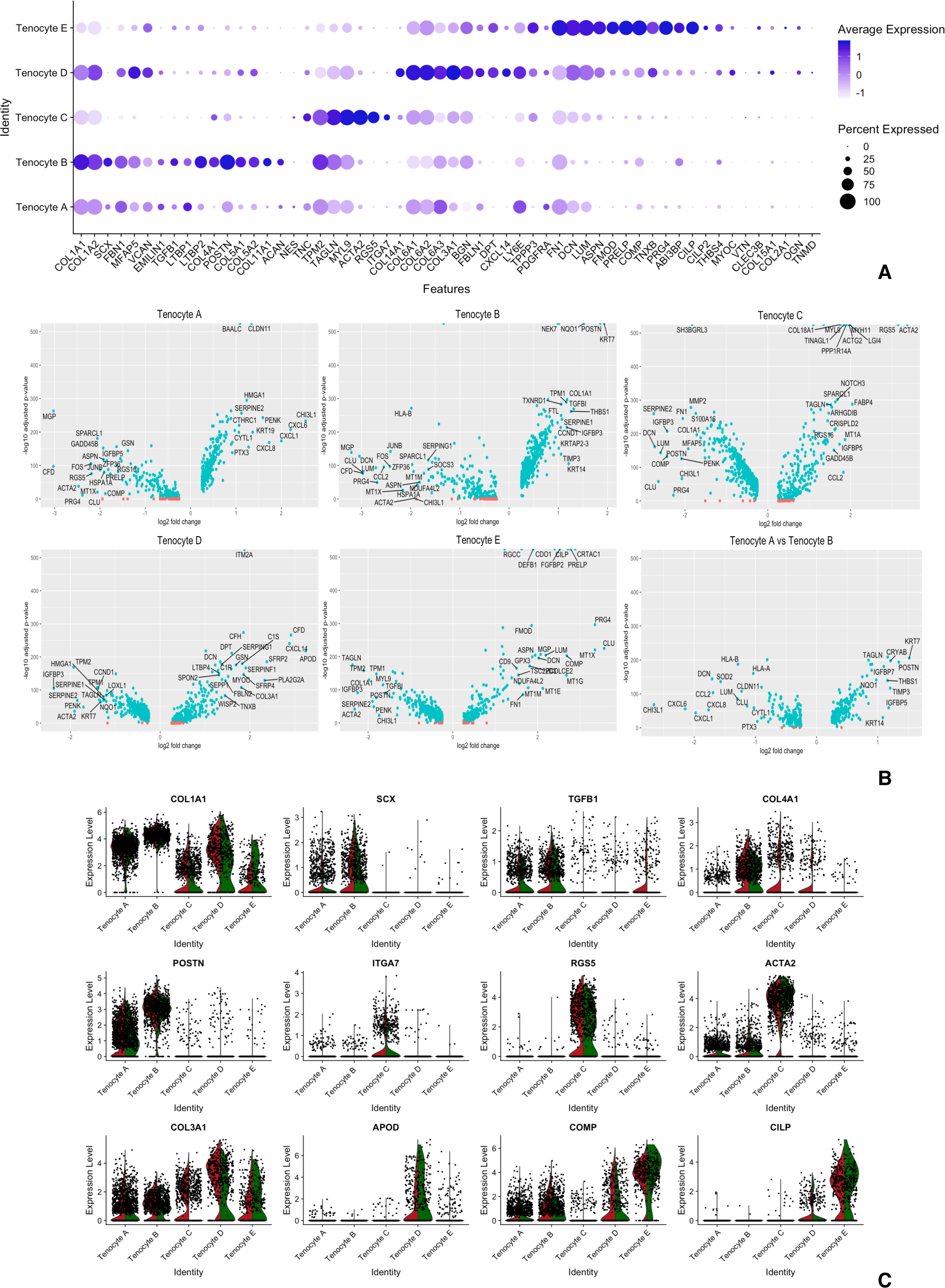
Analysis of five populations of *COL1A* expressing tenocytes. **(A)** Split dot plot of clusters expressing high levels of *COL1A*. The percentage of cells (size of dot) and expression level (intensity of colour) of genes coding for tendon matrix were compared across the five tenocyte populations **(B)** Volcano plots showing differential gene expression of each tenocyte cell clusters compared to the remaining tenocyte populations. **(C)** Split Violin plots of selected matrix genes for diseased (red) versus healthy (green) tendon cells of *COL1A* expressing clusters.

Tenocyte D and Tenocyte E clusters shared features of fibro-adipogenic progenitors (FAPs) including up regulation of *COL3A1, GSN, LUM, DCN, LY6E, PDGFRA* and *CXCL14* (Figure 3). Cells in the Tenocyte D cluster in particular showed up-regulation of *COL6A1/2/3, BGN, FBLN1, APOD, PLA2G2A* and *CXCL14*. Whereas, Tenocyte E cells were found to have increased expression of *PRG4*, *TPPP3*, *DCN, CLU, LUM, PRELP, PCOLCE2, COMP, FMOD, CRTAC1* and *CILP1/2*.

### Single cell surface proteomics reveals a perivascular niche in human tendon

The gene expression signature of cells in the Tenocyte C cluster were characteristic of *ITGA7*+ cells recently described in mouse muscle and named Smooth Muscle-Mesenchymal Cells (SMMCs) (16). These cells have a low expression level of *VCAM1*, and increased expression of *TAGLN, MYL9, ACTA2, RGS5* and *ITGA7* (Figure 3A,C). There was an associated high expression level of basement membrane *COL4A1* (Figure 2B, 3A).

Figure 4 shows the CITE-Seq analysis of oligonucleotide conjugated monoclonal antibodies that recognise surface proteins of the complete disease and healthy tendon data set. SMMCs (Tenocyte C cluster) were found to co-express high levels of surface CD90 and CD146 proteins (Figure 4). In addition, immunohistochemistry demonstrated that ITGA7 positive staining cells were found in human tissue, clustered around vessels (Figure 6F).

**Figure 4.**
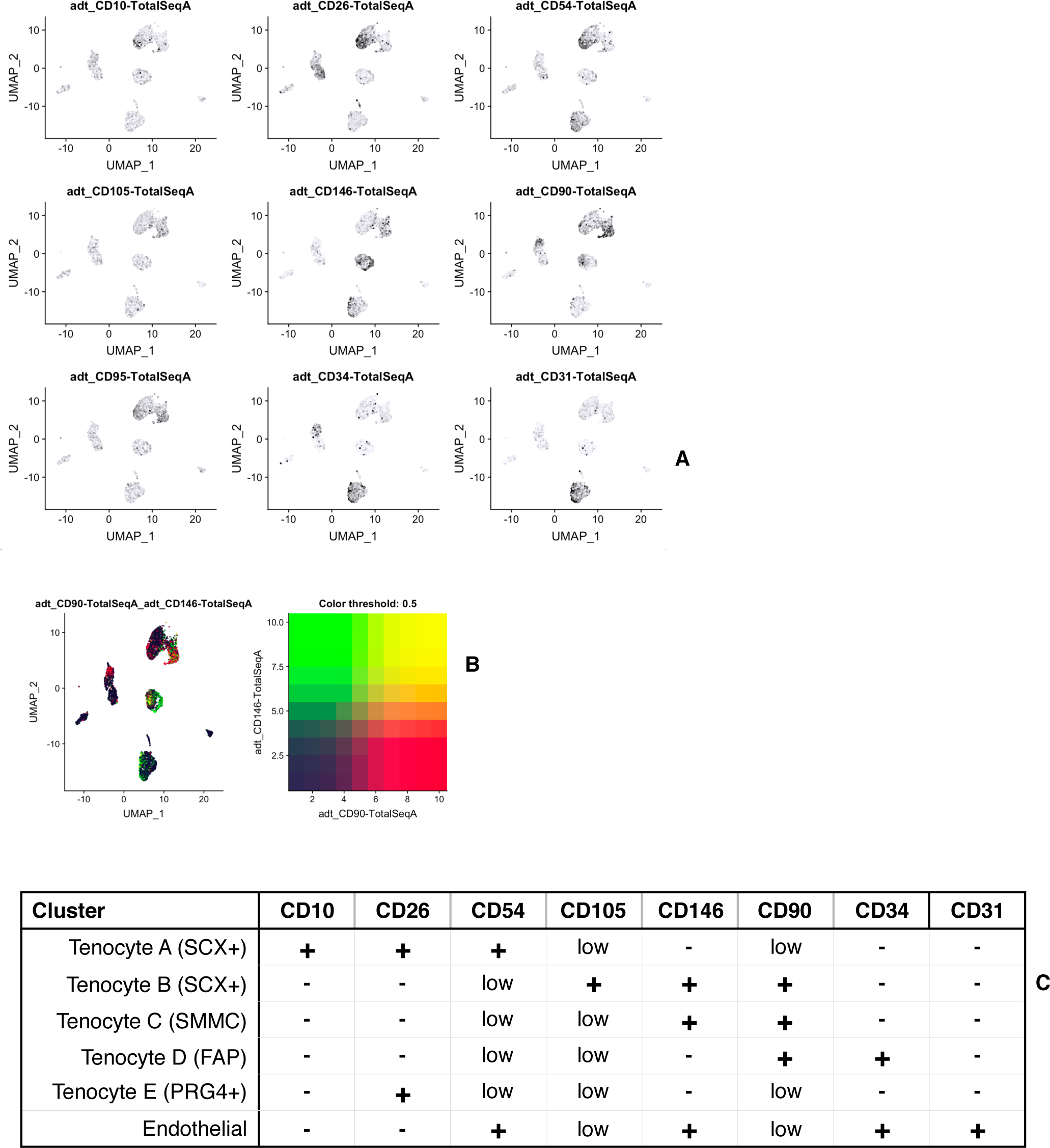
Validation of distinct clusters in human tendon using surface protein quantification. **(A)** Feature plot of ex vivo cells incubated with oligonucleotide barcoded antibodies that recognise surface proteins. **(B)** Combined feature plot demonstrating high co-expression (yellow) of surface CD90 (red) and CD146 (green) on cells in SMMCs of Tenocyte C **(C)** Table summary of differential cell surface markers of non immune cell clusters.

Single cell surface proteomics further distinguished endothelial cells (CD34^+^CD31^+^) from Tenocyte A (CD10^+^CD26^+^CD54^+^/CD90^low^CD95^low^), Tenocyte B cells (CD90^+^CD105^+^CD146^+^), Tenocyte D cells (CD34^+^CD90^+^/CD146^low^CD105^low^) and Tenocyte E cells (CD26^+^/CD90^low^CD54^low^CD10^neg^).

### Stromal cell populations are dynamic in human tendon disease

The Tenocyte A cluster predominantly contained cells from diseased tendon (Figure 2A, S4). This cluster had a similar gene expression profile to Tenocyte B cells but expressed high levels of pro-inflammatory genes including *CXCL1, CXCL6, CXCL8* and *PTX3*. Co-localisation plots shown in Figure 5A illustrate that within the Tenocyte A cluster, the the same cells that expressed *CXCL1* co-expressed *CXCL6* and *CXCL8*. This band of cells predominantly came from diseased tendon samples and very few were found to originate from healthy tendon (Figure 5B, feature plot split by diseased vs healthy tendon). In general, alarmin genes *CD248*, *VCAM1* and *PDPN* were expressed at higher levels by Tenocyte A, B, D and E cells from diseased compared to healthy tendon (Figure 4C). *PTX3* was also up-regulated by diseased cells found in Tenocyte A, Tenocyte B, Tenocyte C, and Tenocyte D. Similarly, there was increased expression of *PDPN, SRRM2, CD248, VCAM1, IL33* and *NES* in endothelial cells from diseased tendon compared to healthy tendon.

**Figure 5.**
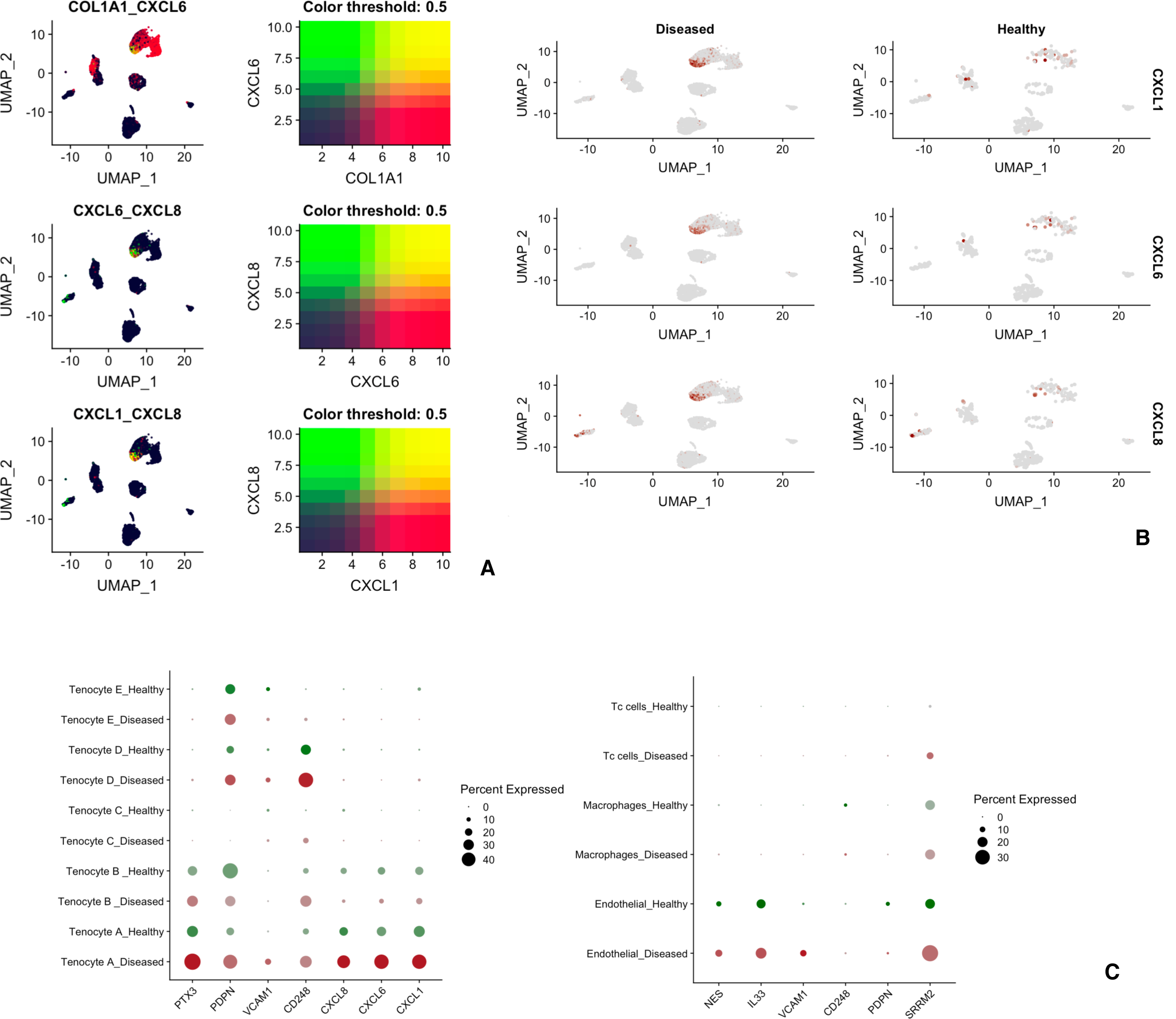
Distinct pro-inflammatory signature of dynamic stromal population in diseased tendon. **(A)** Combined feature plot of ex vivo cells demonstrating co-expression of CXCL6, CXCL1 and CXCL8 by a subset of cells in cluster 0. **(B)** Split feature plot of CXCL6, CXCL1 and CXCL8 by diseased and healthy tendon cells. **(C)** Split dot plot of expression of inflammatory genes by diseased (red) and healthy cells (green) in matrix associated cells (left) and endothelial cells (right). The size of circle indicates percentage of cells expressing the gene and the colour intensity indicates the expression level.

### Discrete cell sub-types are demonstrated in situ in human tissue

Based on the single cell transcriptomic analysis of human tendon cells (Figures 2 and 3), a number of putative protein markers were selected for immuno-histochemistry to identify each of the groups comprising Tenocyte A-E. Four healthy and five diseased tendon samples were formalin fixed and stained with Pentraxin 3 (PTX3) for Tenocyte A, Periostin and Cytokeratin 7 for Tenocyte B, Integrin Subunit Alpha 7 (ITGA7) for Tenocyte C, CXCL14 for Tenocyte D and Matrix gla protein (MGP) for Tenocyte E.

Pentraxin 3 (PTX3) staining demonstrated long linear chains of tendon cells surrounded by matrix (Figure 6C) and these cells were distinct from SMA+CD31+ endothelium (Figure 6A,B). ITGA7 stained cells tended to be situated near blood vessels in small clusters and occasionally formed shorter strings of cells (Figure 6F). Groups of cytokeratin 7 and periostin stained cells were found near the periphery of tendon samples or were clustered within the main substance (Figure 6D, E). Some of these formed chains of cells but without the distinct morphology of PTX3 cells. A minority of cells stained positively for the secreted chemokine CXCL14 and a small group were positive for the intracellular Matrix gla protein (Figure 6H).

**Figure 6.**
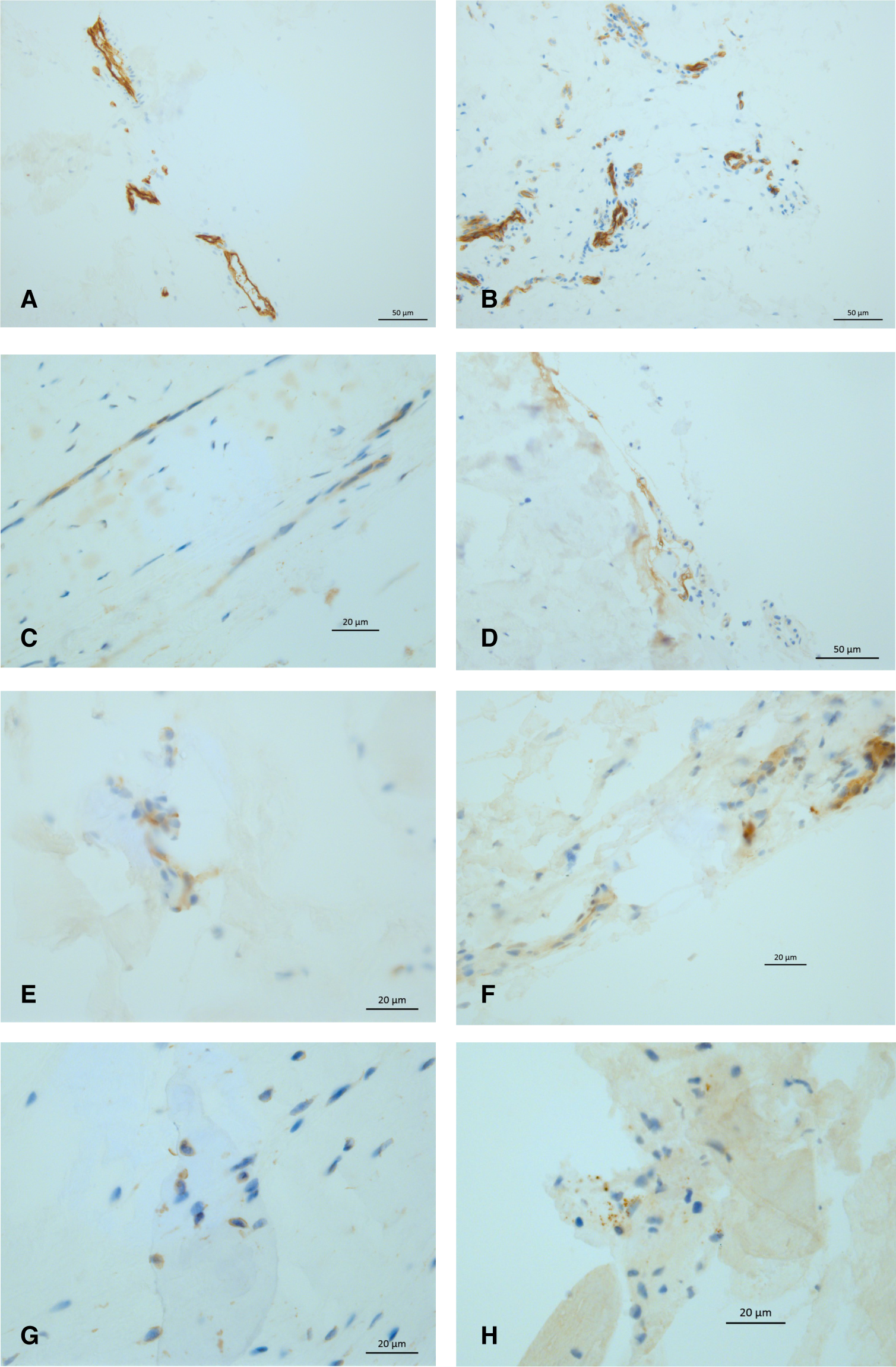
Immunohistochemistry of human tendon demonstrating resident cell sub-types. **(A)** SMA and **(B)** CD31 IHC of endothelial cells. Immuno-staining of human tendon cells with **(C)** PTX3 to identify Tenocyte A; **(D)** periostin and **(E)** Cytokeratin 7 for Tenocyte B; **(F)** ITGA7 for Tenocyte C, **(G)** CXCL14 for Tenocyte D and **(H)** Matrix-gla protein (MGP) for Tenocyte E. The images are representative of four healthy and five diseased human tendon samples.

## Discussion

In the absence of a valid in vivo model of chronic tendon disease, understanding those pathways responsible for tendinopathy has been hampered by conventional methods to isolate crucial cells of interest. In this study, we have combined unbiased single cell gene analysis with surface proteomics of ex vivo single cells and described five sub-populations of *COL1A* expressing human tendon cell (21). In order to highlight the differences between healthy and diseased tissue, samples of three anatomical sites (digital extensor, peroneal and Achilles tendons) were obtained from patients with exclusively end stage tendon disease that had failed to improve despite all non operative measures. To our knowledge this is the first time this approach has been applied to healthy and diseased human tendon.

Five cell clusters were found to express collagen matrix associated genes. The inherent variability in gene expression resulting from technical differences across batches, between different donor individuals and different anatomical sites remains a major limitation of this study (31, 35). Cell clusters were generated using unbiased analysis of differential gene expression but did not include analysis of matrix protein production. It remains possible that the tendon sub-populations identified above represent confounding transcriptomic variation within a single tendon population and/or that a number of cell types failed to survive the digestion process. Hakimi et al. investigated the extracellular proteome of human tendon by comparing healthy and torn shoulder (supraspinatus) tissue (20). As well as identifying a group of proteins up-regulated in diseased tissue, their work serves to provide a catalogue of the most abundant matrix proteins in healthy and diseased human tendon. The gene set of these proteins were therefore used to analyse our data and mapped onto the five discrete *COL1A+* clusters, strengthening the initial unbiased observation of five distinct groups (Figure 3A).

Tenocyte A and Tenocyte B both expressed high levels of *COL1A1/2* and microfibril genes including fibrillin 1 (*FBN1*), versican (*VCAN*), decorin (*DCN*), elastin microfibril interfacer 1 (*EMILIN1*) and microfibril-associated glycoprotein 2/ microfibril associated protein 5 (*MFAP5*). Together these form microfibril chains that have previously been shown to surround a string of tendon resident cells, linking them to the much denser extracellular collagen matrix (11, 22, 23). Histological analysis of human tendon revealed long thin chains of PTX3 positively stained cells that may represent these micro-fibril associated cells (Figure 6C). The observation that microfibrils bind growth factors such as TGFβ and BMP has led to the hypothesis that they play a role in allowing the relatively few resident tendon cells to sample and respond to changes in the surrounding type I collagen matrix (8, 9, 11). Activation of TGFβ signalling involves release of the growth factor from its latent microfibril complex. A number of mechanisms have been described including mechano-transduction, enzymatic degradation, reactive oxygen species and low pH (24). Both *LTBP1* and *LTBP2* expression was greatest in Tenocyte A and Tenocyte B compared to the other matrix cell clusters. *LTBP2* is the only isoform that does not bind to latent TGFβ but rather binds to preformed fibres of fibrillin-1(25). It competes with *LTBP1* for the same binding site and so may provide an additional way in which TGFβ cell signalling is activated in tendon cells by releasing *LTBP1* from microfibrils (26).

Higher levels of *COL4A1, KRT7, POSTN, TAGLN, THBS1, TIMP3, IGFBP5/7* and *COL5A* were seen in Tenocyte B compared to Tenocyte A. It is not clear if Tenocyte B and Tenocyte A represent two separate populations of tendon cells. One of the advantages of CITE-Seq is that it also uses oligo-nucleotide conjugated monoclonal antibodies to recognise cell surface proteins. In this case, Tenocyte B cells were found to have greater levels of surface CD105 and CD146 compared to Tenocyte A cells which were CD10+CD26+CD54+. It remains possible that they share a common progenitor and have developed to perform overlapping roles in response to tissue damage. Alternatively, since Tenocyte A is predominantly composed of diseased tendon cells and virtually all the cells that express chemokine genes *CXCL1, CXCL6* and the cytokine *CXCL8* (IL-8) are from diseased tendon samples, it may be that Tenocyte A cells respond to ECM disruption by recruiting and converting other *COL1A*+ cells to increase their expression of reparative matrix genes.

In addition to *CXCL1, CXCL6* and *CXCL8*, diseased tendon cells in Tenocyte A demonstrate increased expression of inflammatory genes *PDPN* (podoplanin), *VCAM-1* (CD106) and *CD248* (Figure 5). These three were found to be up-regulated in Achilles tendinopathic tissue compared to healthy control tendon (27, 28). We found greatest expression of *VCAM-1, CD248, PDPN* on *SCX*+/microfibril associated cells from Tenocyte A. This further suggests that in the setting of chronic damage, tendon resident *COL1A*+ cells adopt a pro-inflammatory phenotype.

In keeping with previous studies, the predominant immune cell sub-types in tendon are monocytes and Tc cells (28). Similarly, immunostaining of diseased human Achilles samples found reduced levels of IL33 compared to healthy or treated patient samples and this was associated with CD68+ macrophages on immuno-flourescence (29). In our study, *IL33* expression was predominantly found in clusters containing cells with endothelial gene markers and high surface CD31/ CD34, with little expression in *CD68+* clusters or cells expressing collagen genes.

*PTX3* was also found to be increased in diseased cells compared to healthy tendon cells in the Tenocyte A cluster (Figure 5). It encodes a member of the pentraxin protein family and is induced by inflammatory cytokines. *PTX3* has previously been found in endothelial cells and mononuclear phagocytes. In relation to tendon disease, one study found increased *PTX3* in rat model of tendon responses to mechanical stress (30) but there is little prior data on its expression in human tendon disease.

In order to highlight the observed differences between healthy and diseased tendon, the diseased samples were restricted to extreme conditions in which there was end stage tendon disease that remained symptomatic despite all non operative therapies. Our observations are therefore unlikely to be relevant to earlier stages of disease. Furthermore, while there is no significant difference in the male to female distribution of samples in diseased versus healthy samples, diseased samples were obtained from a significantly older patient group (Figure 1). Age related changes in tendon cell gene expression have been found on bulk sequencing (31) and it is possible that this explains the observed differential gene expression between healthy and diseased cells. Finally, a more thorough histological analysis of these inflammatory mediators is required to appreciate their relevance in tendon pathophysiology.

Previous single cell transcriptomic studies of mouse muscle have demonstrated a number of different cell matrix associated cell populations including fibro-adipogenic progenitors (FAPs), muscle satellite cells (MuSCs), *ITGA7+VCAM1-*/Smooth Muscle-Mesenchymal Cells (SMMCs), and scleraxis (*SCX)* expressing cells (16) (32) (14) (15). Giordani et al identified up to 8 different cell clusters found in mouse muscle (16). One of these groups was designated FAPs based on expression of *LY6A, LY6E, PDGFRA* and *DCN*. Increased expression of *LY6E, PDGFRA* and *DCN*, but not *LY6A*, was found in cells in both Tenocyte D and Tenocyte E from both diseased and healthy human tendon cells (Figure 2, 3) raising the possibility that these are also types of FAPs. In comparison to Tenocyte E, cells in Tenocyte D exhibited greater expression of tendon collagen genes *COL1A1/2, COL3A1;* microfibril genes *FBN1* and *MFAP5*; and chemo-attractants *CXCL2, CXCL14.* Whereas, Tenocyte E cells expressed *TPPP3* and *PRG4* found in putative tendon stem cells that reside in the paratenon of mice patella tendon and respond to acute tendon injury by producing reparative matrix (17). In our cohort, these cells co-expressed genes associated with cartilage formation including *ASPN, COMP, PCOLCE2, FMOD, CILP, FN1* and *PRELP* and were largely found in clusters around disorganised matrix (Figure 6). Further work is required to explore whether these differences represent a genuine phenotypic deviation in which reparative *PRG4+* cells switch to produce a more cartilaginous matrix in the chronic setting; perhaps as a last ditch effort to maintain some structural integrity.

Smooth Muscle-Mesenchymal Cells (SMMCs) were defined in mouse muscle by Giordani et al. as a group of *VCAM*-cells that express *ITGA7, RGS5* and *MYL9* (16). When mouse SMMCs were isolated, a myogenic subset were found to promote muscle growth, forming chimeric myotubes in culture with muscle satellite cells (MuSCs). We found a similar cluster of cells in both healthy and disease human tendon (Figure 2, 3) and theses tended to be situated around vessels (Figure 6). To our knowledge this is the first demonstration of these cells in human tendon and, in keeping with their original description, they do not express endothelial markers found in pericytes (33). Proteomic analysis revealed surface expression of CD90 and CD146 on these human cells (Figure 5).

One explanation for the presence of SMMCs may be that tendon samples were taken close the myo-tendinous junction and so muscle tissue was inadvertently included. While this may be a possibility for the healthy hamstring tendon samples, even despite of careful dissection, the SMMCs cluster consisted of cells from all tendon samples including the majority taken well distal to the muscle insertion. The tendon-bone enthesis interface was also avoided and this is supported by the low expression of *COL2A1* genes across all the clusters (Figure 3A). Interestingly, *SCX* positive ‘tendon fibroblasts’, similar to Tenocyte B and Tenocyte A, were found in mouse muscle devoid of tendon tissue by Giordani et al. Further work could investigate whether these two seemingly out of place cells are the remnants of respective populations that once had a developmental role or if they still have an adult function, for example a co-ordinated structural response to injury of both muscle and tendon.

One previous study of human tendon used Fluidigm single cell analysis to identify a group of nestin+ putative tendon stem/progenitor cells. This was limited to 71 human tendon cells, all cultured to passage 1 and analysed using 46 gene transcripts. It identified three main cell clusters including a minority expressing increased levels of *NES/CD31/CD146*((34). In our data set of over 6,000 cells obtained immediately ex vivo from human tendon, *NES* (nestin) expression was predominantly found in endothelial cell cultures co-expressing *PECAM-*1, *CD34* and surface CD34/CD31 (Figure 2,3 and S5). This fits with the published nestin+ immunostaining of human Achilles tendon that coincided with blood vessel endothelium. A small percentage of endothelial cells in our study co-expressed *COL1A1* but there was little co-expression of other tendon cell markers such as *SCX* (Figure S6). While subsequent mouse models of tendon development and healing demonstrated a role for nestin+ cells, further work is required to demonstrate the importance of these endothelial associated cells in adult human tendon. It is possible that by culturing cells to passage 1, collagen producing nestin+ cells were preferentially selected by Yin et al. In comparison to ex-vivo cells, we identified eight clusters of in vitro cultured cells from three tendon samples, the smallest of which expressed endothelial markers and contained nestin+ cells (Figure S6). All eight clusters contained cells expressing increased levels of *COL1A1* and a minority co-expressed *NES*. While there were clusters recognisable as Tenocyte A, B, C and D, cells expressing SMMC markers were not evident. For these reasons we excluded passaged cells from further analysis.

## Conclusion

Comparative integrated analysis of the transcriptome of human tendon cells obtained ex vivo from healthy and diseased samples revealed multiple sub-populations expressing matrix associated genes. These included FAP type cells, clusters co-expressing *SCX* and microfibril associated genes, and the first demonstration of *ITGA7*+ Smooth Muscle-Mesenchymal Cells (SMMCs) and *TPPP3/PRG4+* cells in adult human tendon. CITE-Seq surface proteomics identified surface markers by which these cells could be isolated in future. *SCX+* microfibril gene expressing cells from diseased tendon expressed high levels of pro-inflammatory markers including *CXCL1, CXCL6, CXCL8, PDPN* and *PTX*, and diseased endothelium up-regulated *NES, IL33* and *PDPN*.

### Special thanks to

Hubert Slawinski, Theo Kyriakou, Santiago Revale and Rory Bowden (Oxford Genomics Centre) and Molly Browne and Leticia Campo (Department or Oncology, Oxford MSD)

---

**Supplemental Figure 1.**
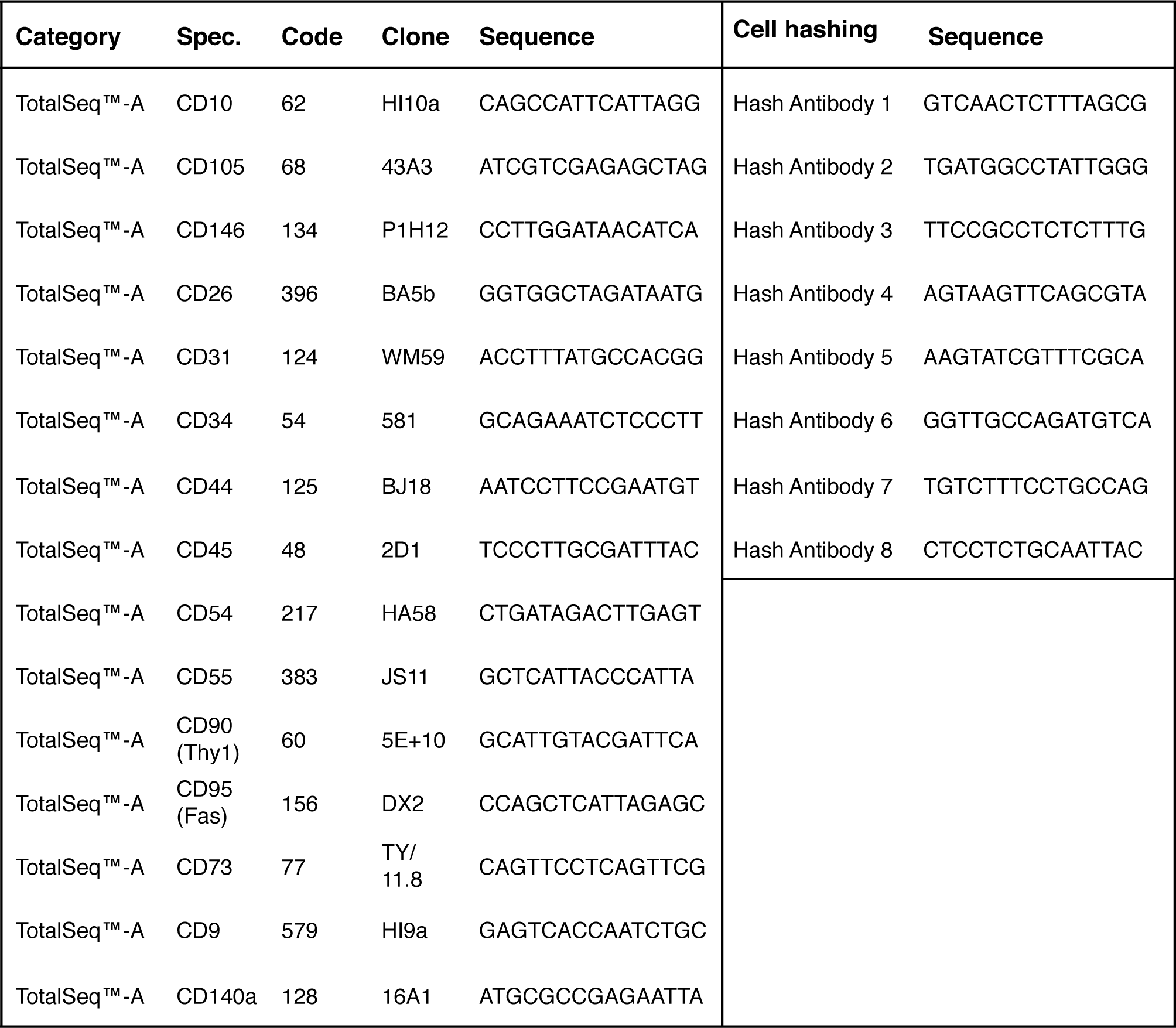
CITE-Seq utilises oligonucleotide barcodes conjugated to monoclonal antibodies against cell surface proteins. Fifteen candidate cell surface proteins were targeted. A further eight hashing antibodies, that recognise ubiquitous surface proteins, were exploited to identify cells from one of eight different tendon samples run on a single lane of sequencing.

**Supplemental Figure 3.**
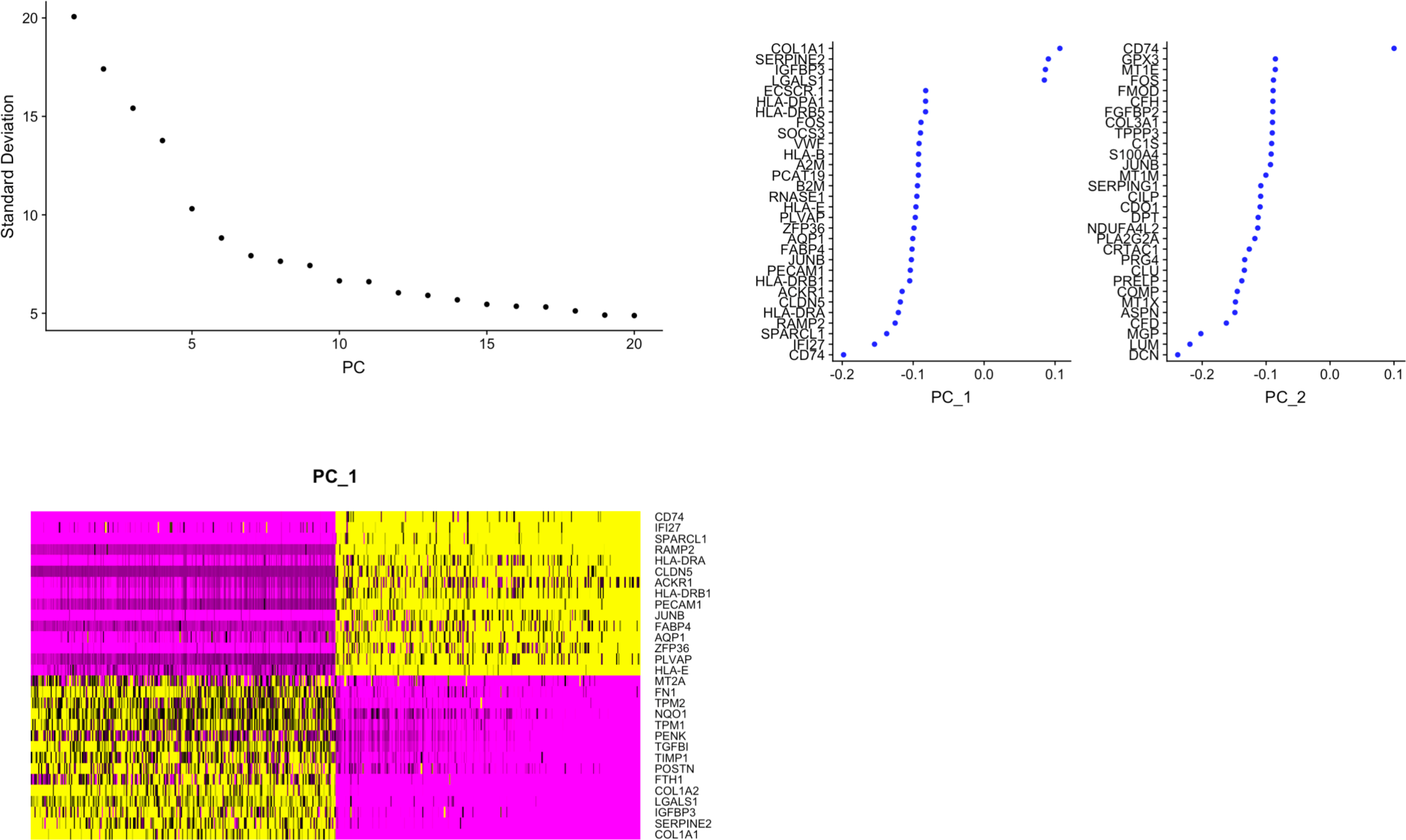
Principal component analysis and Elbow plot of the integrated ex vivo diseased and healthy data set following quality control.

**Supplemental Figure 2.**
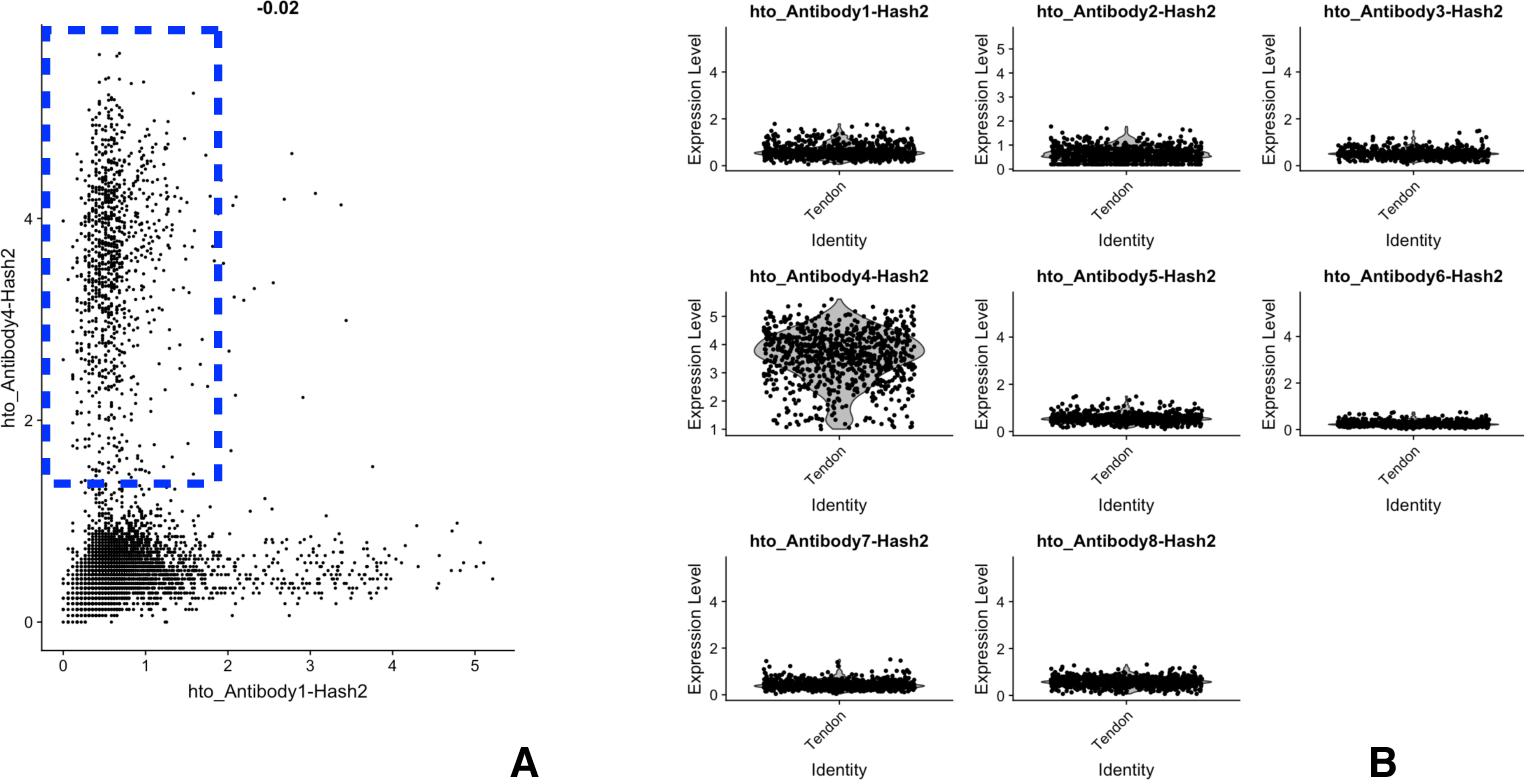
Cells from a given tendon sample were incubated with CITE-Seq monoclonal antibodies conjugated to one of eight oligonucleotide barcodes (hash mAb) that bind ubiquitous surface proteins. Post sequencing, the relative ‘expression level’ of these hash barcode mAb were used to identify all the cells from a particular tendon sample. This meant that cells from up to eight tendon samples could be sequenced in a single lane. In this example, Seurat v3 was used to select all cells with high ‘expression level’ of hashing hto_Antibody4 and low ‘expression level’ of hto_Antibody1 (**A**). Subsequent Violin plots of the selected cell data showed low ‘expression level’ of the other seven hashing antibodies, so confirming the purity of the selected population. This process was performed for all samples before they were integrated as a combined diseased versus healthy data set.

**Supplemental Figure 4.**
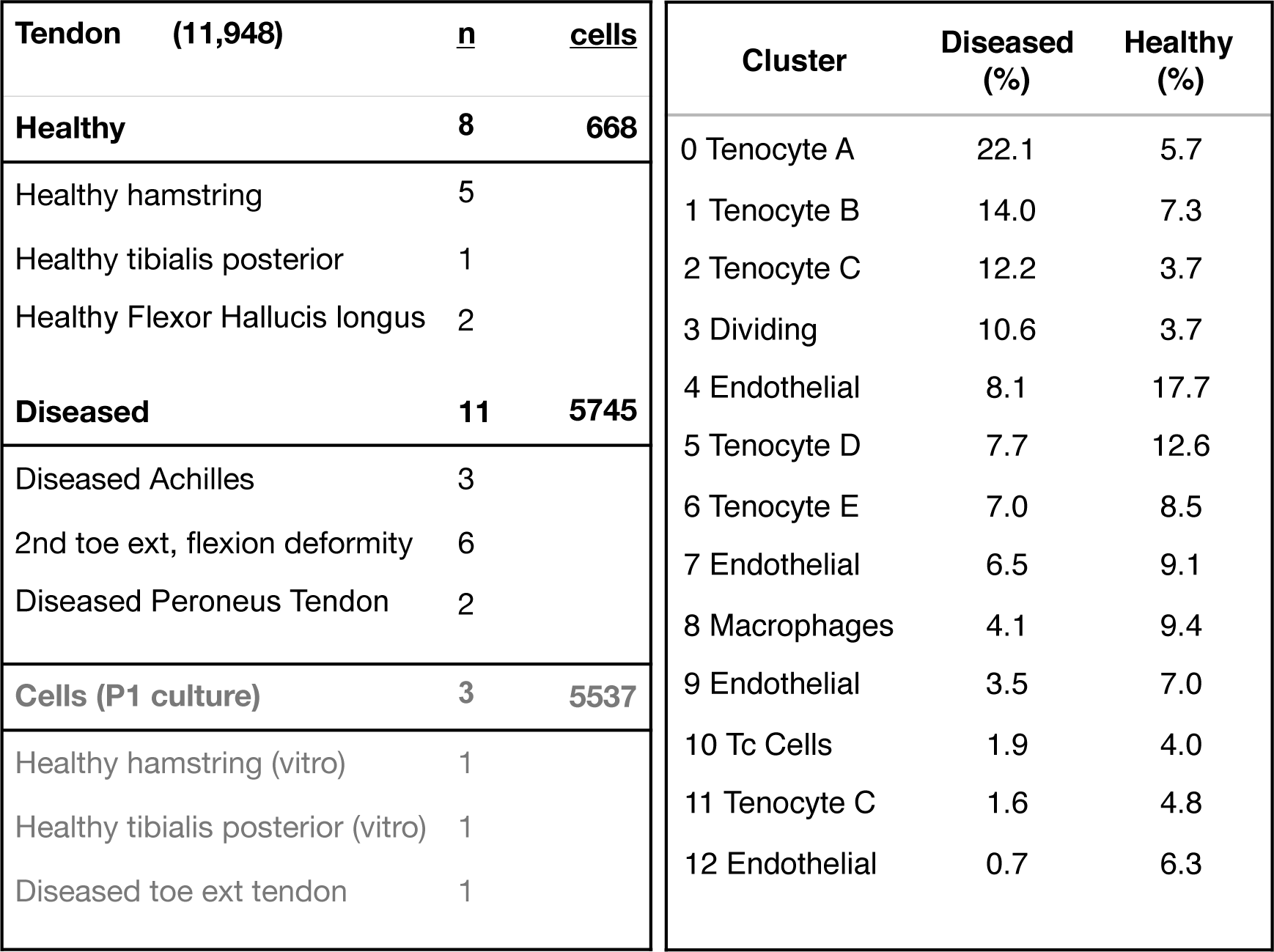
Cells were obtained from three main sources; ex vivo healthy human tendon, ex vivo diseased tendon and cells cultures to passage 1 from two healthy and one diseased tendon. A total of 11,948 cells were analysed using Seurat V3 post quality control. The right hand table quantifies the distribution of the integrated ex vivo data set across the clusters identified by Seurat v3.

**Supplementary figure 5.**
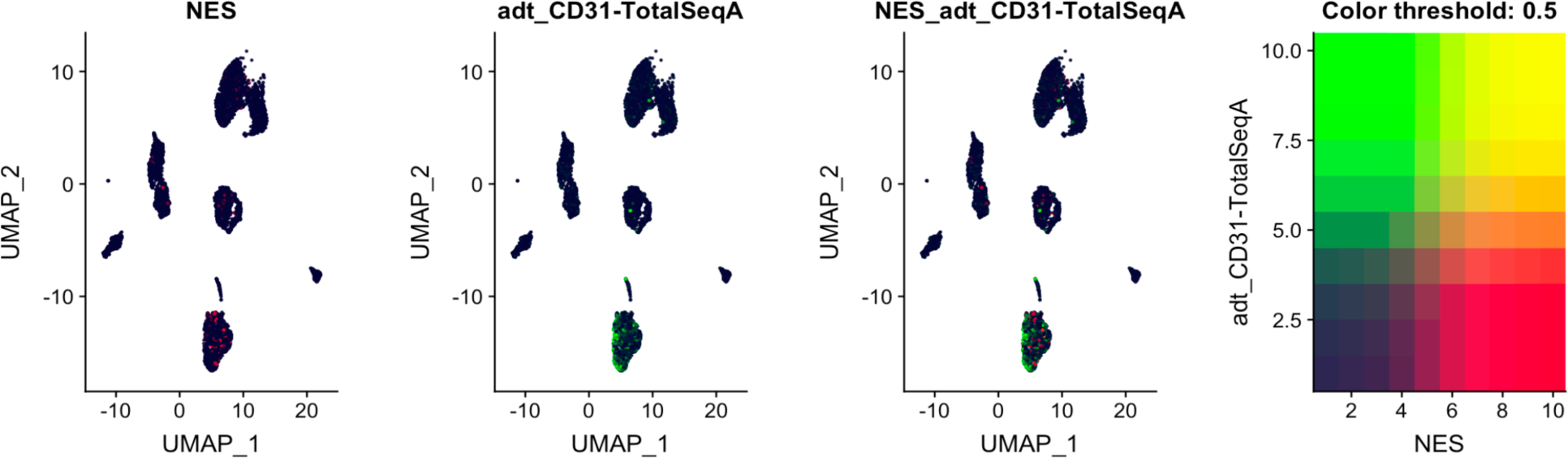
CITE-Seq combined feature plot of ex vivo cells demonstrating co-expression of *NES* gene and surface CD31 on cells within clusters expressing endothelial gene markers.

**Supplemental Figure 6.**
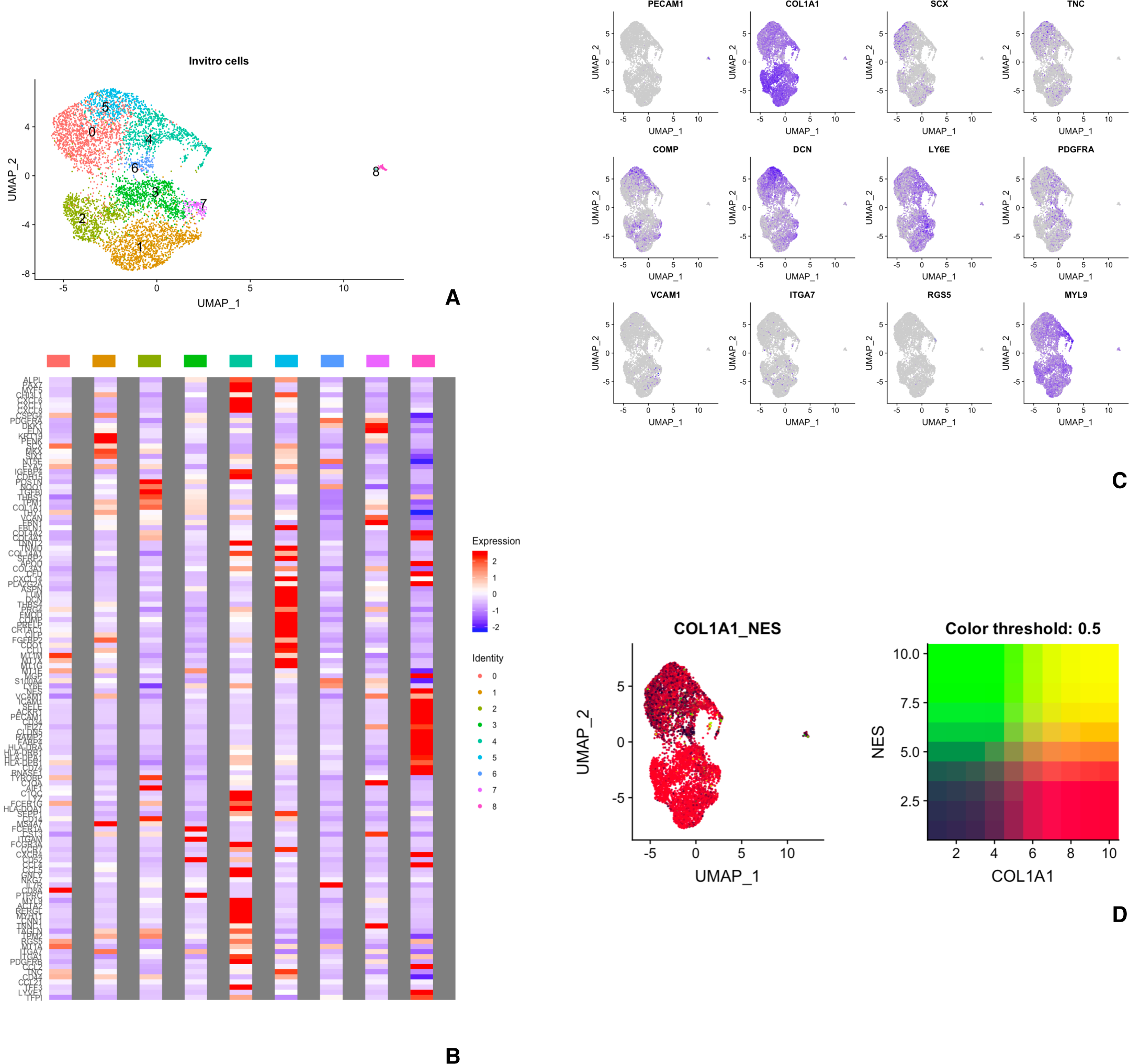
Integrated CITE-Seq analysis of three tendon samples (two healthy and one diseased) cultured in vitro until passage 1 following mechanical and enzymatic dissociation. **(A)** UMAP demonstrating eight clusters. **(B)** Heat map of average gene expression across the eight clusters of the same gene set for ex vivo cells (see Figure 2). **(C)** Feature plot of selected canonical markers to help identify clusters. **(D)** Combined feature plot of in vitro cultured cells co-expressing NES and COL1A1.

